# Multiple origin but single domestication led to *Oryza sativa*

**DOI:** 10.1101/127688

**Authors:** Jae Young Choi, Michael D. Purugganan

## Abstract

The domestication scenario that led to Asian rice (*Oryza sativa*) is a contentious topic. Here, we have reanalyzed a previously published large-scale wild and domesticated rice dataset, which were also analyzed by two studies but resulted in two contrasting domestication models. We suggest the analysis of false positive selective sweep regions and phylogenetic analysis of concatenated genomic regions may have been the sources that contributed to the different results. In the end, our result indicates Asian rice originated from multiple wild progenitor subpopulations; however, *de novo* domestication appears to have occurred only once and the domestication alleles were transferred between rice subpopulation through introgression.

## Introduction

Asian rice (*Oryza sativa*) is a diverse crop comprising of five subpopulations (Garris *et al*. 2005). Archaeobotanical data for japonica and indica, the two major subpopulation of *O. sativa*, suggests japonica was domesticated first ~7000 years ago in the Yangtze Basin of China while indica was domesticated later ~4000 years ago in the Ganges plains of India (Fuller *et al*. 2010). Elucidating the origins of Asian rice and the history of its domestication has been a contentious field (Gross and Zhao 2014). With whole genome data, it is becoming apparent that each Asian rice variety group/subspecies (aus, indica, and japonica) had distinct subpopulations of wild rice (O. *nivara* or *O. rufipogon*) as its progenitor (Huang *et al*. 2012). Specifically, wild rice could be divided into three major subpopulations and were designated as Or-I, Or-II, and Or-III by Huang *et al*. (2012). Phylogenetically, japonica was most closely related to Or-III, while aus and indica were most closely related to Or-I but each was monophyletic with a different subset of Or-I samples (Huang *et al*. 2012). These results were consistent with a recent whole genome study showing distinct wild progenitors led to aus, indica, and japonica domestication (Choi *et al*. 2017).

Whether rice was domesticated once and subsequent varieties were formed by introgression with different wild progenitors, or whether each variety was domesticated independently in different parts of Asia, is debatable. The debate mainly arose from two studies analyzing the same data but surprisingly arriving at two different domestication scenarios: Huang *et al*. (2012) supporting the single domestication with introgression model with Civáň *et al*. (2015) supporting the multiple domestication model. If the causal domestication mutation arose in a single genotype and was subsequently introgressed into the other two subpopulations (single domestication with introgression model), gene trees for the domestication region would differ from the genome-wide tree. Because the domestication is hypothesized to have occurred once in a single subpopulation and spread to the other subpopulations through hybridization, we define it as the single *de novo* domestication with introgression model. On the other hand, if the domestication mutation arose in each subpopulation on a different genetic background and was independently selected (multiple domestication model), then the gene trees for the domestication region would be concordant with the genome-wide tree. As domestication should leave evidence of recent selection, both studies used a reduction in polymorphism levels as a metric to detect local genomic regions associated with domestication. The evolutionary histories of those regions were then interpreted as the domestication history for Asian rice.

We argue there may have been two issues that may have led to the conflicting results between Huang *et al*. (2012) and Civáň *et al*. (2015). First, regions that were detected by the two studies are candidate selective sweep regions that may or may not be related to domestication. Even population genetic model-based methods of detecting selective sweeps are prone to false-positives and with the right condition any evolutionary scenario can be interpreted with a false-positive selective sweep region (Pavlidis *et al*. 2012). Because lineage sorting within a genomic region depends on the effective population size (N_e_) of that region (Pamilo and Nei 1988), evolutionary factors that decreases N_e_ (e.g. population bottleneck, positive or negative selection, and low recombination rate) can accelerate the lineage sorting process (Hobolth *et al*. 2011; Scally *et al*. 2012; Prüfer *et al*. 2012; Pease and Hahn 2013). Given that each Asian rice had separate wild progenitor population of origin, any false-positive selective sweep region will likely to be concordant with the underlying species phylogeny, and spuriously support the multiple domestication model. Hence, any candidate domestication related selective sweep region would need additional evidence before being considered for downstream evolutionary analysis. Second, both studies used genotype calls made from a low coverage (1~2X) resequencing data (Huang *et al*. 2012). Uncertainty associated with genotype calls made from low coverage data (Nielsen *et al*. 2011) could be another source that led to the different results for the two studies.

Thus, we revisited the domestication scenarios proposed by the two studies and reanalyzed the Huang *et al*. (2012) data using a complete probabilistic framework that takes the uncertainty in SNP and genotype likelihoods into consideration (Fumagalli *et al*. 2014; Korneliussen *et al*. 2014). We then carefully compared our results against the two domestication models and contrasted it against the results from both Huang *et al*. (2012) and Civáň *et al*. (2015) studies.

## Materials and Method

Raw paired-end FASTQ data from the Huang *et al*. (2012) study was download from the National Center for Biotechnology Information website under bioproject ID numbers PRJEB2052, PRJEB2578, PRJEB2829. We excluded the aromatic rice group from the analysis, as their sample sizes were too small and we excluded the few samples that had too high coverage. In the end a total of 1477 samples were selected for analysis (Supplemental Table 1).

Raw reads were then trimmed for adapter contamination and low quality bases using trimmomatic ver. 0.36 (Bolger *et al*. 2014) with the command:

~~~
java –jar trimmomatic-0.36.jar PE \
        $FASTQ1 $FASTQ2 \
        $FASTQl_paired $FASTQl_unpaired $FASTQ2_paired $FASTQ2_unpaired \
         ILLUMINACLIP:TruSeq2-PE.fa:2:30:10:4 \
         LEADING:3 TRAILING:3 SLIDINGWIND0W:4:15 MINLEN:30
~~~

Quality controlled FASTQ reads were then realigned to the reference japonica genome downloaded from EnsemblPlants release 30 (ftp://ftp.ensemblgenomes.org/pub/plants/). Reads were then mapped to the reference genome using the program BWA-MEM ver. 0.7.15 (Li 2013) with default parameters. Alignment files were then processed with PICARD ver. 2.9.0 (http://broadinstitute.github.io/picard/) and GATK ver. 3.7 (McKenna *et al*. 2010) toolkits to remove PCR duplicates and realign around INDEL regions (DePristo *et al*. 2011).

Using the processed alignment files, genotype probabilities were calculated with the program ANGSD ver. 0.913 (Korneliussen *et al*. 2014). The genotype probabilities were then used by the program ngsTools (Fumagalli *et al*. 2014) to conduct population genetic analysis. To estimate theta (θ) ngsTools uses the site frequency spectrum as a prior to calculate allele frequency probabilities. Usually site frequency spectrum requires an appropriate outgroup sequence to infer the ancestral state of each site. However, for calculating Watterson and Tajima’s θ it is not necessary to know whether each polymorphic site is a high or low frequency variant (Korneliussen *et al*. 2013). Hence, we used the same reference japonica genome as the outgroup but strictly for purposes of calculating θ. Per site allele frequency likelihood was calculated using ANGSD with the commands:

~~~
angsd –b $BAMLIST –ref $REF –anc $REF –out $SFS –r $CHR \
    –uniqueOnly 1 –remove_bads 1 –only_proper_pairs 1 -trim 0 \
    –C 50 –baq 1 –minMapQ 20 –minQ 30 \
    –minlnd $minlnd \
    –setMinDepth $setMinDepth \
    –setMaxDepth $setMaxDepth \
    –doCounts 1 -GL 1 -doSaf 1
~~~

Per site allele frequency for each domesticated and wild subpopulation was calculated separately with different filtering parameters using the options –minInd, –setMinDepth, –setMaxDepth; where parameter minInd represent the minimum number of individuals per site to be analyzed, setMinDepth represent minimum total sequencing depth per site to be analyzed, setMaxDepth represent maximum total sequencing depth per site to be analyzed. Specifically, -minInd and –setMinDepth were set as a third of the number individuals in the subpopulation while –setMaxDepth was set as five times the number individuals in the subpopulation. Overall site frequency spectrum was then calculated with the realSFS program from the ANGSD package. Using each subpopulation’s site frequency spectrum as prior, we then calculated θ for each subpopulation using ANGSD with the command:

~~~
angsd –b $BAMLIST –ref $REF –anc $REF –out $THETA –r $CHR \
    –uniqueOnly 1 –remove_bads 1 –only_proper_pairs 1 –trim 0 \
    –C 50 –baq 1 –minMapQ 20 –minQ 30 –minlnd $minlnd \
    –setMinDepth $setMinDepth \
    –-setMaxDepth $setMaxDepth
    –doCounts 1 –GL 1 –doSaf 1 \
    –doThetas 1 –pest $SFS
~~~

Using the output file from the previous command, for each subpopulation a sliding window analysis was then conducted with the thetaStat program from the ANGSD package using non-overlapping window length and step sizes of 20 kbp, 100 kbp, 500 kbp, and 1000 kbp with the command:

~~~
thetaStat do_stat $ALLELEFREQ_POSTPROB_FILE \
      –nChr $IND \
      –win $WINDOW –step $STEP
~~~

For each window, θ per site was estimated by dividing Tajima’s theta (θ_π_) against the total number of sites with data in the window. Windows with less then 25% of sites with data were discarded from downstream analysis. This resulted in a minimum of 90% of the windows being analyzed (Supplemental Table 2). Sweeps were identified using sliding windows that were estimating the ratio of wild to domesticate polymorphism (π_w_/π_d_). To calculate π_w_/π_d_ values we chose the Or-II subpopulation to calculate π_w_ since Or-II subpopulation was most distantly related to all three domesticated rice subpopulation (Supplemental Fig 1). πw/πd values were calculated separately for each domesticated rice subpopulations. Windows with large π_w_/π_d_ values were designated as candidate domestication selective sweep region, and significance was determined using an empirical distribution of π_w_/π_d_ values described below.

Japonica has demographic history that is consistent with more intense domestication related bottlenecks then aus and indica (Xu *et al*. 2011). Thus, many π_w_/π_d_ values for japonica are expected to be similar between true domestication sweep and neutral regions, causing difficulties in identifying true-positive selective sweeps. Hence, we chose the approach of Civáň *et al*. (2015) by using a single threshold π_w_/π_d_ value to determine significance for all three subpopulation. In contrast to Civáň *et al*. (2015) we chose our threshold based on the empirical distribution of each subpopulation. The 97.5 percentile π_w_/π_d_ values were determined for each domesticated rice subpopulation, and the subpopulation with the lowest 97.5 percentile π_w_/π_d_ values was decided as the significance threshold. The threshold percentile that is represented by each subpopulation and window size is listed in Supplemental Table 3. If the top 2.5% windows in the subpopulation with the lowest π_w_/π_d_ threshold are all due to true domestication sweeps, then in the other two subpopulations the threshold may be seen after a selective sweep or a population bottleneck. These co-located low-diversity genomic regions (CLDGRs) then represent candidate domestication-related selective sweep regions for all three subpopulations, and it is necessary for each CLDGR to have additional information to differentiate itself from the background domestication-related bottleneck scenarios. We assumed CLDGRs overlapping genes with functional genetic evidence related to domestication phenotypes (Meyer and Purugganan 2013) as true candidate domestication genes. Custom R (R Core Team 2016) code for identifying the overlapping selective sweep windows can be found in the GitHub repository: https://github.com/cjy8709/Huangetal2012_reanalysis.

To account for the uncertainty in the underlying data, phylogenetic analysis were conducted by estimating pairwise genetic distances from genotype probabilities (Vieira *et al*. 2016). We ran the program ANGSD to calculate genotype probabilities for all 1477 domesticated and wild rice samples using the command:

~~~
angsd –b $BAMLIST –ref $REF –out $GEN0PP –r $CHR \
    –uniqueOnly 1 –remove_bads 1 –only_proper_pairs 1 –trim 0 \
    –C 50 –baq 1 –minMapQ 20 –minQ 30 \
    –minlnd $minlnd
    –setMinDepth $setMinDepth
    –setMaxDepth $setMaxDepth
    –doCounts 1 –GL 1 –doMajorMinor 1 –doMaf 1 \
    –skipTriallelic 1 –SNP_pval le-3 –doGeno 8 –doPost 1
~~~

Initially, the effects of different filtering parameters on the downstream phylogenetic analysis were examined by using three different parameter values for the options – minInd, -setMinDepth, -setMaxDepth: 1) minInd=492, setMinDepth=492, setMaxDepth=4920; 2) minInd=738, setMinDepth=738, setMaxDepth=8862; 3) minInd=492, setMinDepth=369, setMaxDepth=8862. Afterwards all subsequent phylogenetic analysis were conducted with genotype posterior probabilities calculated using the minInd=492, setMinDepth=492, setMaxDepth=4920 parameter set. Genotype posterior probabilities were then used by the program ngsDist from the ngsTools package to estimate all pairwise genetic distances. Using the output file from the previous command, neighbor-joining trees were reconstructed with the genetic distances using the program FastME ver. 2.1.5 (Lefort *et al*. 2015) with the command:

~~~
fastme–2.1.5-linux64 –D 1 –i $DISTANCE_FILE –o $TREE
~~~

Phylogenetic trees were diagramed using the web interface iTOL ver. 3.4.3 (http://itol.embl.de/) (Letunic and Bork 2016).

## Results and Discussions

In both Huang *et al*. (2012) and Civáň *et al*. (2015) studies, the phylogeny based on genome-wide data versus putative domestication regions were compared to determine which domestication scenario was best supported by the data. We reconstructed the genome-wide phylogeny by estimating genetic distances between domesticated and wild rice using genotype probabilities (Vieira *et al*. 2016). To examine the effects of different parameters would have on downstream analysis, three different parameters were used to estimate the genotype probabilities. The probabilities were then subsequently used to estimate genetic distances and build neighbor-joining trees for each chromosome (Supplemental Fig 1). Comparing trees built from the three different parameters, each chromosomal phylogeny was largely concordant with the others forming two major clades where the japonicas were grouping together, while indica and aus formed a monophyletic group (Figure 1A). Further, the trees corroborated the results of Huang *et al*. (2012) where the japonicas were most closely related to the Or-III wild rice subpopulation, while indica and aus were most closely related to the Or-I wild rice subpopulation.

**Figure 1.**
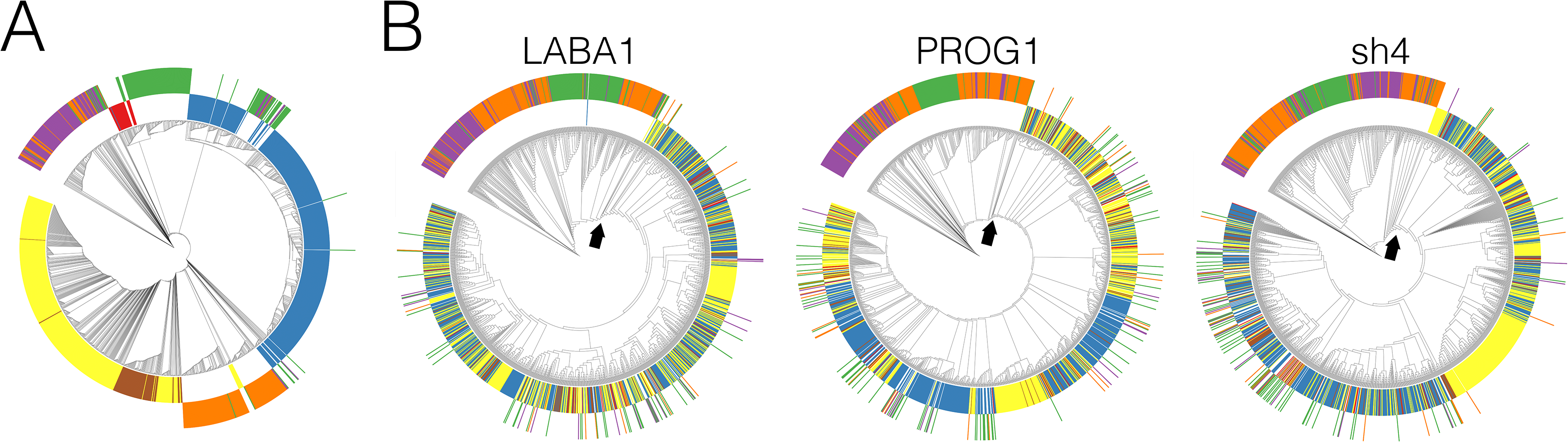
Neighbor-joining tree for A) chromosome 1 and B) 20 kbp upstream and downstream (40 kbp total) of domestication genes *LABA1, PROG1*, and *sh4*. Inner circle of colors represent domesticated rice: red, aus; blue, indica; yellow, temperate japonica; brown, tropical japonica. Outer circle of colors represent wild rice that were designated by Huang *et al*. (2012): green, Or-I; purple, Or-II; orange, Or-III. Arrows indicate the most ancestral internal node of the monophyletic domesticated rice clade.

We then scanned for local genomic regions associated with domestication related selective sweeps to infer the domestication history of Asian rice. Compared to the π_w_/π_d_ method, there are population genetic model based tests that are more powerful for detecting selective sweeps (Vitti *et al*. 2013). However, these methods use hard called genotypes and do not take the uncertainty associated with low coverage data into account. But more importantly the two previous study of Huang *et al*. (2012) and Civáň *et al*. (2015) used the π_w_/π_d_ method to detect domestication related sweeps. For consistency we also implemented the w/ d method using genotype likelihoods to take the low coverage into consideration, and took a closer look at the genomic regions with significant evidence of a selective sweep.

To identify putative selective sweep regions, we chose the approach of Civáň et al. (2015) and identified sweep regions separately for each rice subpopulation. If rice had a single *de novo* domestication of origin in one subpopulation and the other two subpopulations were domesticated through introgression, then all three rice subpopulations would have identical sweep regions with shared haplotypes; otherwise, the single *de novo* domestication with introgression model cannot be supported. CLDGRs (Civáň *et al*. 2015) were identified using a window size that was different from both Huang *et al*. (2012) (100 kbp) and Civáň *et al*. (2015) (between 100 and 200 kbp), using a 20 kbp sliding window to narrow down on the candidate genes relating to domestication. To identify significant CLDGRs we chose a stringent cutoff to conservatively identify candidate regions and identified a total of 39 CLDGRs (Supplemental Table 4).

Neighbor-joining trees were then reconstructed for each 39 CLDGRs (Supplemental Figure 2). The majority of CLDGRs showed monophyletic relationships among the domesticated rice subpopulation, where japonica, indica, and aus were clustering between and not within subpopulation types. Six windows (e.g. chr2:11,660,000-11,680,000) showed phylogenetic relationships where each domesticated sample were clustering with the same domesticated subpopulation type.

This initially suggested the evolutionary history of CLDGRs were most consistent with the single *de novo* domestication model. We then examined larger window sizes of 100 kbp, 500 kbp, and 1000 kbp for candidate CLDGRs (Supplemental Table 4) and reconstructed phylogenies for those regions (Supplemental Fig 3,4, and 5). Larger window sizes have fewer numbers of windows for analysis, hence leading to fewer numbers of CLDGRs being identified (Supplemental Table 2). Nonetheless, with increasing window sizes CLDGR phylogenies became more congruent with the genome-wide phylogenies, consistent with the multiple domestication model. However, CLDGRs are only candidate regions that may harbor domestication genes or may be false positive selective sweep regions affected by domestication-related bottlenecks. As population bottlenecking can decrease effective population sizes, false positive CLDGRs may represent regions of the genome with increased lineage sorting. These regions are then likely to have phylogenies that are more concordant with the underlying species phylogeny (Pamilo and Nei 1988). Hence, it is crucial that a CLDGR have additional evidence that can associate it with selection and differentiate its evolutionary history from the underlying species phylogeny. To do so we searched CLDGRs that overlapped genes with functional genetic evidence related to domestication. We found three known domestication genes: long and barbed awn gene *LABA1* (chr4:25,959,399-25,963,504), the prostrate growth gene *PROG1* (chr7:2,839,194-2,840,089), and shattering locus *sh4* (chr4:34,231,186-34,233,221) (Li *et al*. 2006; Tan *et al*. 2008; Hua *et al*. 2015). Interestingly, the gene *sh4* was the only gene detected across multiple sliding window sizes excluding the largest 1000 kbp window (Supplemental Table 4).

Phylogenetic trees were then reconstructed for the three domestication loci that included 20 kbp upstream and downstream (40 kbp in total) of their coding sequence. We note for all three genes the casual variant resulting in the domestication phenotype were located in the protein coding sequences (Li *et al*. 2006; Jin *et al*. 2008; Hua *et al*. 2015). For all three genomic regions, the phylogenies were clustering different subpopulation types of domesticated rice together (Figure 1B), consistent with the single *de novo* domestication scenario.

Interestingly, *sh4* was identified as a candidate gene with evidence of selective sweep in this study and both Huang *et al*. (2012) and Civáň *et al*. (2015). Only Civáň *et al*. (2015) did not find evidence of single origin in a phylogenetic tree reconstructed from a 240 kbp region surrounding *sh4*. A single nucleotide mutation in the *sh4* gene causes a reduction in seed shattering (Li *et al*. 2006), which is an important domestication trait thought to minimize the labor during harvesting (Sang and Ge 2007). Sanger sequencing of the *sh4* region across various domesticated rice have indicated a near identical haplotype at the *sh4* region suggesting a single origin of non-shattering (Li *et al*. 2006; Zhang *et al*. 2009; Thurber *et al*. 2010). When we reconstructed phylogenies for 40 kbp windows surrounding the *sh4* region, the upstream region of the start codon had phylogenies where the domesticated rice were clustering with the same domesticated subpopulation types (Supplemental Fig 6). We then reconstructed the phylogeny for large genetic regions surrounding each three domestication loci and discovered with each increased window size, the phylogeny of the region increasingly corroborated the genome-wide phylogeny by clustering with the same subpopulation type (Figure 2). This was not only true for *sh4*, but also *LABA1* and *PROG1* as well. Thus, the domestication-related evolutionary history for *sh4* is limited to the gene and regions that are downstream of the stop codon. Including large flanking regions or concatenating candidate domestication region without functional genetic evidence can lead to phylogenies that are concordant with the genome-wide species phylogeny, spuriously concluding it as evidence for the multiple domestication origin model. In fact, careful re-examination of the Huang *et al*. (2012) results indicate their phylogeny for the concatenated 55 candidate domestication region actually shows a topology where japonica and indica were grouping with the same domesticated subpopulation type, while individual domestication regions with functional evidence showed a mix of japonica and indica clustering together, suggesting the concatenation had resulted in the neutral diversity of the non-domestication related regions to overwhelm the domestication regions phylogenetic signatures.

**Figure 2.**
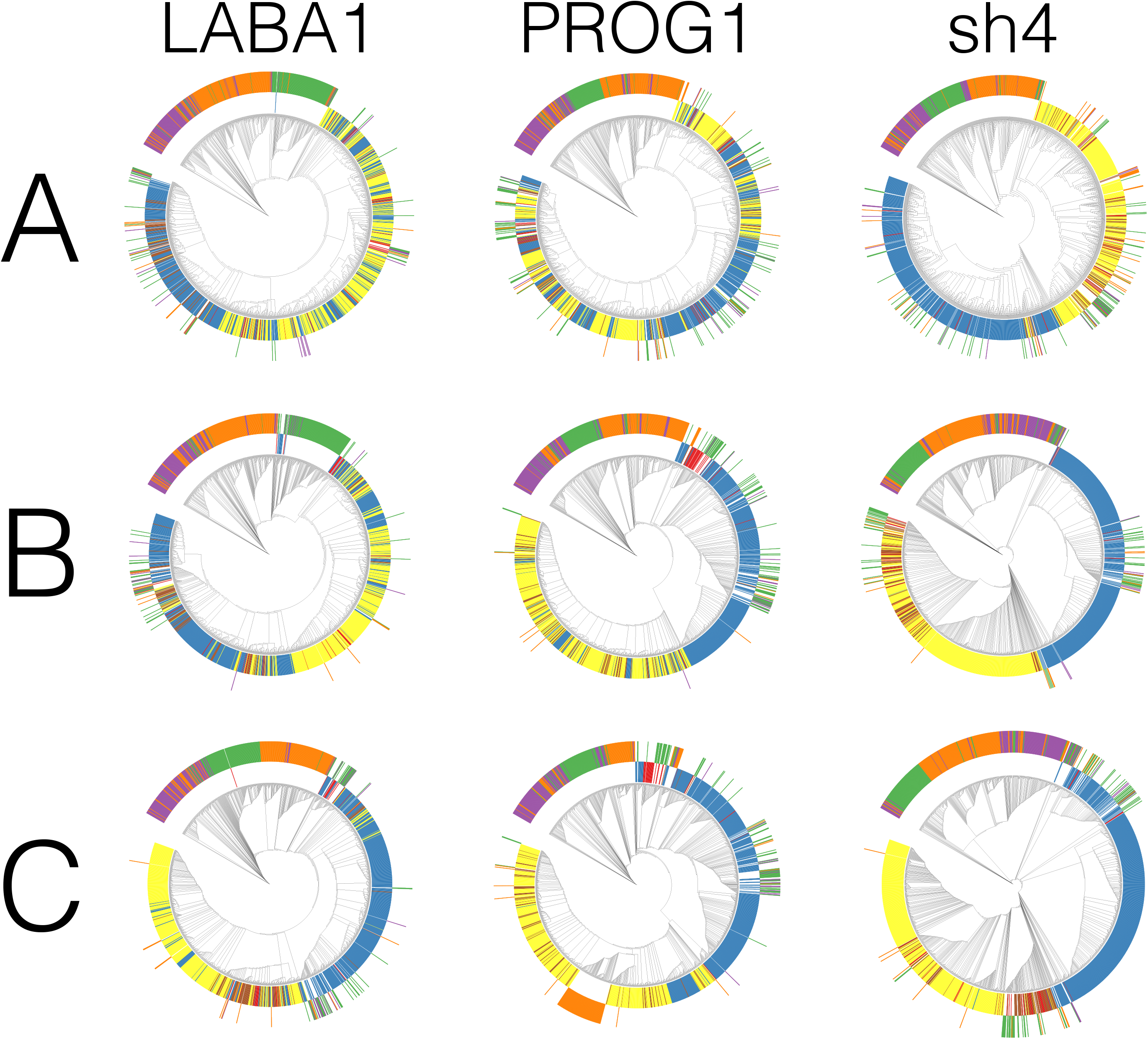
Neighbor-joining trees for three different window sizes flanking the domestication genes *LABA1, PROG1*, and *sh4*. First row, 50 kbp upstream and downstream of gene (100 kbp total); Second row, 250 kbp upstream and downstream of gene (500 kbp total); Third row, 500 kbp upstream and downstream of gene (1000 kbp total).

In this study we have used the same approach as Huang *et al*. (2012) and Civáň *et al*. (2015) to search for regions of domestication related selective sweeps and investigated those regions’ evolutionary history. With stringent thresholds and conservative assumptions to exclude false positive CLDGRs we were able to narrow down to three genes (*LABA1, PROG1*, and *sh4*), which were likely to be the key genes involved in the domestication of Asian rice (Meyer and Purugganan 2013). We note that our method of detecting selective sweeps may have missed other true domestication sweeps when other methods are applied (Liu *et al*. 2017), and the three genes may represent the minimum number of genes involved in the domestication of Asian rice. With higher coverage data becoming available (Li *et al*. 2014) powerful population genetic model based selective sweep test can be applied to detect more loci that were potentially involved in the domestication process. But even with these methods one is still left with candidate regions and it is likely that different studies would detect different regions with significant evidence of undergoing a selective sweep, which could potentially result in contrasting domestication scenarios between studies. Hence, it is important that future domestication studies using advanced methods for detecting selection should consider whether the candidate region would also have functional genetic evidence as well.

Civáň *et al*. (2015) had criticized the role of *PROG1* and *sh4* in domestication due to several wild rice alleles clustering with the domesticated alleles (Figure 1B). However, evidence from de-domesticated weedy rice shows feralized rice can carry the causative domestication allele but not retain any of the domestication phenotypes (Li *et al*. 2017), suggesting some of the wild rice in the Huang et al. (2012) dataset may actually represent different stages of feralized domesticated rice (Wang *et al*. 2017). Further, Civáň *et al*. (2015) claimed the clustering of wild and domesticated rice alleles as evidence of selection from standing variation (i.e. soft sweep), which led to the observed phylogenies for *PROG1* and *sh4* (Civáň and Brown 2017). However, given the deep genome-wide divergence between japonica and indica subpopulation (Choi *et al*. 2017; Wang *et al*. 2017), soft sweeps in this case is expected to produce distinct haplotypes and not homogenize the haplotypes between subpopulations (Messer and Petrov 2013). Thus, clustering of wild rice with domesticated rice in candidate domestication genes is more consistent with the frequent gene flow occurring between domesticated and wild rice (Wang *et al*. 2017). Subsequently, we caution the interpretation of phylogeographic analyses investigating the geographic localities of wild and domesticated CLDGRs, because phylogenetic clustering between wild and domesticated alleles can not differentiate originating progenitor vs. recent hybridization between wild and domesticated rice.

Recently Wang *et al*. (2017) suggested several wild rice samples in the Huang *et al*. (2012) study had evidence of gene flow from domesticated rice into the wild rice. Thus, the genetic affinity between Or-I wild rice subpopulation and indica/aus; and Or-III wild rice subpopulation and japonica, may represent recent gene flow and it is unclear whether Or-I and Or-III wild rice subpopulation represents the direct progenitor of domesticated rice. The deep genome-wide coalescence time between japonica and indica predating the archaeologically estimated domestication time (Choi *et al*. 2017) suggests japonica and indica has independent wild rice of origin, but its possible this progenitor was not sampled in the Huang *et al*. (2012) study. This is clearly seen across the genome-wide phylogeny of the domesticated rice as all samples cluster with the same domesticated subpopulation type. On the other hand, phylogenies from domestication loci were consistent with a single origin model where all domesticated subpopulations were monophyletic with each other. Further, in all three regions the most closely related wild rice corresponded to the Or-III subpopulation, supporting the hypothesis that the domestication alleles were introgressed from japonica into indica and aus (Huang *et al*. 2012; Choi *et al*. 2017). There is a possibility that the Or-III subpopulation positioning as the sister group to all domesticated rice in domestication-associated genomic regions may be an artifact from the gene flow between japonica and wild rice (Wang *et al*. 2017). However, we do not think this is the case for three reasons: 1) gene flow from domesticated to wild rice is predominately originating from indica and aus subpopulations (Wang *et al*. 2017), 2) gene flow of domestication allele into wild rice will cluster wild and domestication rice together not positioning them as a separate sister group, and 3) even if it is a result of gene flow the sister group position suggests it was an old gene flow event ultimately involving the japonica subpopulation.

In the end, our evolutionary analysis for the domestication loci *LABA1, PROG1*, and *sh4* are consistent with both Sanger and next-generation sequencing results (Li *et al*. 2006; Tan *et al*. 2008; Xu *et al*. 2011; Huang *et al*. 2012; Hua *et al*. 2015). Our results are also consistent with the archaeological and genomic evidences (Fuller *et al*. 2010; Choi *et al*. 2017). Here then, we provide support for a model in which Asian rice has evolved from multiple origins but *de novo* domestication had only occurred once (Caicedo *et al*. 2007; Gao and Innan 2008; He *et al*. 2011; Huang *et al*. 2012; Castillo *et al*. 2016; Choi *et al*. 2017) (Figure 3). Specifically, this model hypothesizes each domesticated rice subpopulation had distinct wild rice subpopulation as its immediate progenitor, but *de novo* domestication only occurred once in japonica, involving genes that include *LABA1, PROG1*, and *sh4*. The domestication alleles for these genes were then subsequently introgressed into the wild progenitors of aus and indica by gene flow and ultimately led to their domestication. Indeed crossing between wild and domesticated rice has been common practice in modern rice breeding for enhancing the limited domesticated rice genetic pool (Brar and Khush 1997), and may have been the ultimate source of diversifying the initial proto-domesticated rice into genetically differentiated domesticated rice subpopulations.

**Figure 3.**
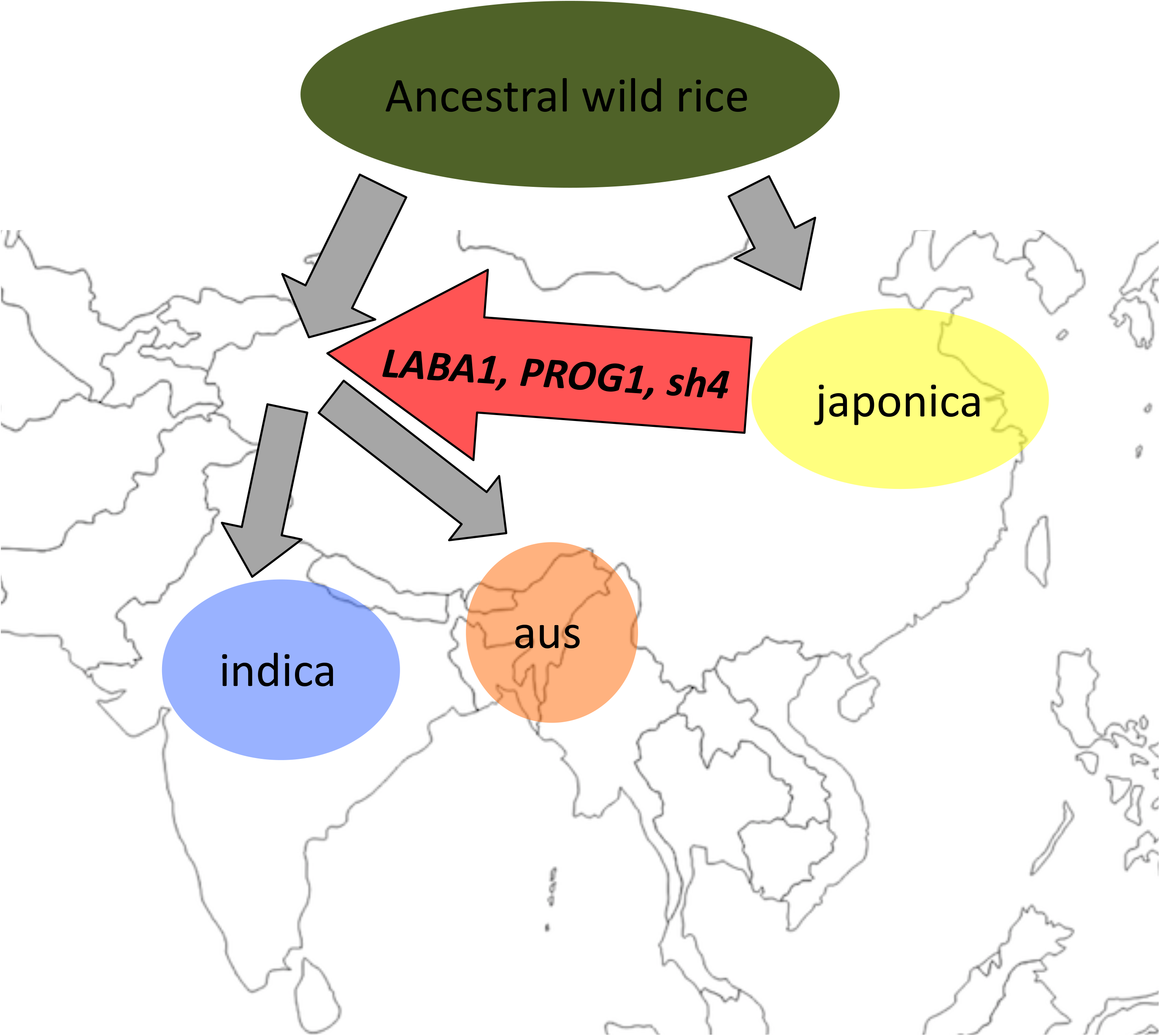
Domestication scenario that led to Asian rice. Each domesticated rice subpopulation had separate wild rice progenitor, however because the geographic origin of the progenitor is heavily debated its location is omitted from the map. Geographic position of the domesticated rice represent hypothesized domestication areas and was based on Fuller *et al*. (2010). Red arrow indicates a gene flow event that transferred the japonica originating domestication haplotypes from the genes *LABA1, PROG1*, and *sh4* into the progenitors of aus and indica, which led to their domestication.

## Acknowledgements

This work was supported by grants from the National Science Foundation Plant Genome Research Program (IOS-1546218), the Zegar Family Foundation (A16-0051) and the NYU Abu Dhabi Research Institute (G1205) to M.D.P. We appreciate the New York University – High Performance Computing for providing computational resources and support. We thank Simon “Niels” Groen, Zoe Lye, and the anonymous reviewers for providing helpful comments.

---

Supplemental Fig1. Neighbor-joining trees for 12 chromosomes. Each row represent trees built using genotype posterior probabilities calculated from 3 different parameters. Top, middle, and bottom row represents tree built from genotype posterior probabilities calculated from parameter 1, 2, and 3; listed in materials and method.

Supplemental Fig2. Neighbor-joining trees for the 39 CLDGRs identified after 20 kbp sliding window.

Supplemental Fig3. Neighbor-joining trees for the 10 CLDGRs identified after 100 kbp sliding window.

Supplemental Fig4. Neighbor-joining trees for the 4 CLDGRs identified after 500 kbp sliding window.

Supplemental Fig5. Neighbor-joining trees for the 2 CLDGRs identified after 1000 kbp sliding window.

Supplemental Fig6. Neighbor-joining trees for 40 kbp windows surrounding the gene sh4 (chr4:34,231,186-34,233,221).

